# Healthy soils, rising pests? Stopping plowing improves soil quality but increases wireworm infestation: evidence from a long-term field study

**DOI:** 10.64898/2025.12.16.694608

**Authors:** Le Cointe Ronan, Plantegenest Manuel, Morvan Thierry, Kevin Levardois, Menasseri Safya, Poggi Sylvain

**Affiliations:** IGEPP, INRAE, Institut Agro, Univ Rennes, 35653, Le Rheu, France; IGEPP, INRAE, Institut Agro, Univ Rennes, 35000, Rennes, France; INRAE, UMR1069 Sol Agro et hydrosystème Spatialisation, F-35000 Rennes, France

**Keywords:** Plowing, soil health, wireworms, bulk density, aggregate stability, soil moisture, pest dynamics, sustainable farming

## Abstract

Conventional tillage (i.e. plowing) is often associated with soil degradation and a loss of biodiversity. In response, reduced tillage has increasingly been promoted as a sustainable alternative for its positive impacts on soil structure, water dynamics, and biodiversity. However, reduced tillage not only favors beneficial soil-dwelling organisms, but also pests. In this field study, we examined the effects of five years of reduced tillage on the physical properties of the soil and the dynamics of *Agriotes* wireworm populations (Elateridae), which are increasingly widespread pests. Our results show that tillage reduction led to stable edaphic conditions and a redistribution of organic matter, creating a favorable environment for wireworms. While our results demonstrate the beneficial effects of reduced tillage on aggregate stability, they also indicate a concomitant increase in soil bulk density, suggesting a reduction in water-holding capacity. Monitoring of wireworm populations revealed their aggregated distribution and their increase in abundance in infested areas year on year. Monitoring soil moisture revealed that tillage reduction improved water dynamics, enhancing infiltration and reducing evaporation. This could potentially favor the development of wireworms. Surprisingly, wireworm size distribution showed a higher proportion of young instars in plowed plots, evidencing that, firstly, the lack of soil cover does not prevent oviposition and that injury caused by plowing targets more the last instars rather than young larvae. While reduced tillage improves key soil health indicators, our findings suggest a potential trade-off in terms of increased pest pressure. Our study highlights the importance of adopting a holistic approach when designing sustainable cropping systems, as well as considering the services and dis-services they provide.

## Introduction

Designing sustainable cropping systems requires an understanding of the services and dis-services they provide. Reduced-tillage (RT) farming is widely recognized for its positive impacts on soil structure, water dynamics, and biodiversity (Henneron et al., 2015; Holland, 2004; Kladivko, 2001; Power, 2010), while being economically very competitive (Verch et al., 2009). These benefits are further enhanced when reduced tillage is combined with organic amendments, which contribute to improved soil fertility and biological activity (Annabi et al., 2011; Bottinelli et al., 2013; Edmeades, 2003). While the benefits of reduced tillage on soil are well-documented (Skaalsveen et al., 2019) the balance between beneficial and detrimental soil functions it provides (Power, 2010) occasionally diminishes the overall agronomic and ecological gains (Brennan et al., 2014; Palm et al., 2014). For instance, RT improves water infiltration in the upper soil layers and increases soil organic matter content (Rasmussen, 1999), but it can impair nutrient cycling (Brennan et al., 2014). Moreover, reduced tillage often leads to an increase in soil bulk density (SBD), particularly in silty soils, which can reduce macroporosity. In addition, RT influences not only the soil organic carbon distribution (Moreno et al., 2006), physical properties of the soil but also, directly or indirectly, its biological functioning, specifically when combined with organic amendment (Bottinelli et al., 2017). The effect of edaphic conditions on soil organisms is complex, underlining the importance of adopting a holistic approach when designing cropping systems. Crotty et al. (2016) further illustrated this complexity by showing that tillage can simultaneously reduce earthworm abundance while promoting certain nematode populations, highlighting the trade-offs involved in tillage decisions.

Wireworms, the subterraneous larvae of click beetles (Coleoptera: Elateridae: *Agriotes*), are increasingly widespread pests in Europe (Poggi et al., 2021). In France, wireworms are the most damaging pests on maize and potatoes. In Brittany, western France, in 2021 damage was recorded in 14% of potato plots, leading to €2.5 million in direct costs for farmers. As wireworms have a multi-year life cycle (2–5 years), with generations overlapping, they can accumulate over the crop rotation, depending on agro-environmental factors. Wireworms thrive in undisturbed soils, which provide them with a constantly moist and protected habitat ideal for their development (Le Cointe et al., 2023). By contrast, plowing causes them direct mechanical injury or exposes them to bird predation, desiccation, and unfavorable conditions. However, wireworms often inhabit soil layers deeper than those affected by tillage, providing them some refuge from mechanical disturbance. Wireworm populations exhibit seasonal vertical migration, with their peak presence in the upper soil layers occurring in spring and autumn (Furlan, 1998; Gratwick M, 1989) regulated by soil moisture and temperature conditions (Jung et al., 2014; Lafrance, 1968). We hypothesize that vertical migration is less pronounced in environments with more stable, buffered edaphic conditions.

The present study aims at evaluating the effects of plowing versus reduced tillage (i.e., rotary harrow) combined with two fertilization treatment on edaphic conditions, as well as their influence on wireworm population dynamics. Five years after the beginning of a split-plot experiment, the minimum duration required for system stabilization (Bai et al., 2018), with tillage practices as main plots and fertilization regimes as subplots, we monitored key soil parameters including the vertical distribution of soil organic carbon, soil bulk density, aggregate stability, and soil moisture variability, concurrently with wireworm (genus *Agriotes*) abundance and spatiotemporal distribution over four consecutive years. This comprehensive approach allowed for an integrated assessment of the effects of tillage on the abiotic and biotic components of soil health.

## Materials and Methods

### Study site and crop rotation

The experimental site was the EFELE experimental platform (INRAE UMR SAS) located at Le Rheu in western France (48°06′07 N, 1°47′44 W). The local climate is temperate, with an average annual temperature and annual cumulative precipitation of 12 °C and 650 mm, respectively. The soil type is Luvic Cambisol to Haplic Luvisol (Association française pour l’étude du sol, 2008) with a silt loam texture (15% clay, 69% silt, 16% sand in the A-horizon). The experiment started in 2012 on a field cultivated in maize during the previous two years and mouldboard-plowed to 25 cm before the experiment. Initial soil water pH-value was 5.1 and soil organic carbon content was 20.6 g/kg of soil in the top layer (0-30 cm). The experimental crop rotation consists in a two-year succession of maize (*Zea mays L.*) and winter wheat (*Triticum aestivum L*.) with white mustard (*Sinapis alba*) as a nitrate-catch cover crop between the wheat and the maize. Wireworm monitoring began in 2017 in winter wheat, after five years of experimentation.

### Experimental designs

#### A split-plot design with tillage practices as main plots and fertilizer treatments as subplots

The factorial experimental design combined two tillage practices and two nitrogen fertilizer treatments resulting in four treatments replicated three times. The 12 plots were arranged in a split-plot design, with tillage practices as main plots and fertilizer treatments as subplots (see Fig.1). Each individual plot was 32 m long and 12 m wide. The first tillage treatment (referred thereafter as conventional tillage, CT) consisted in moldboard plowing with full soil inversion to a depth of 25 cm, followed by harrowing to 12 cm before seeding. The soil was plowed in spring for maize and in autumn for wheat. The second tillage treatment (referred thereafter as reduced tillage, RT) consisted in soil tillage with a rotating harrow, without soil inversion, to a depth of 12 cm. Nitrogen fertilizer treatments consisted in the application of ammonium nitrate every year for mineral treatment (MF) and the application of cattle manure every two years (50t/ha) before maize sowing supplemented with mineral fertilizers on wheat for organic treatment (OF). Nitrogen inputs were applied at once on maize before sowing and twice on winter wheat at tillering and heading stages. Nitrogen fertilizer doses were calculated according to a projection of nitrogen balance and to an analysis of the applied manure (dry weight, nitrogen content, carbon content). On average, 90 kg/ha of nitrogen were applied on maize and 120 kg/ha on winter wheat.

**Fig. 1.**
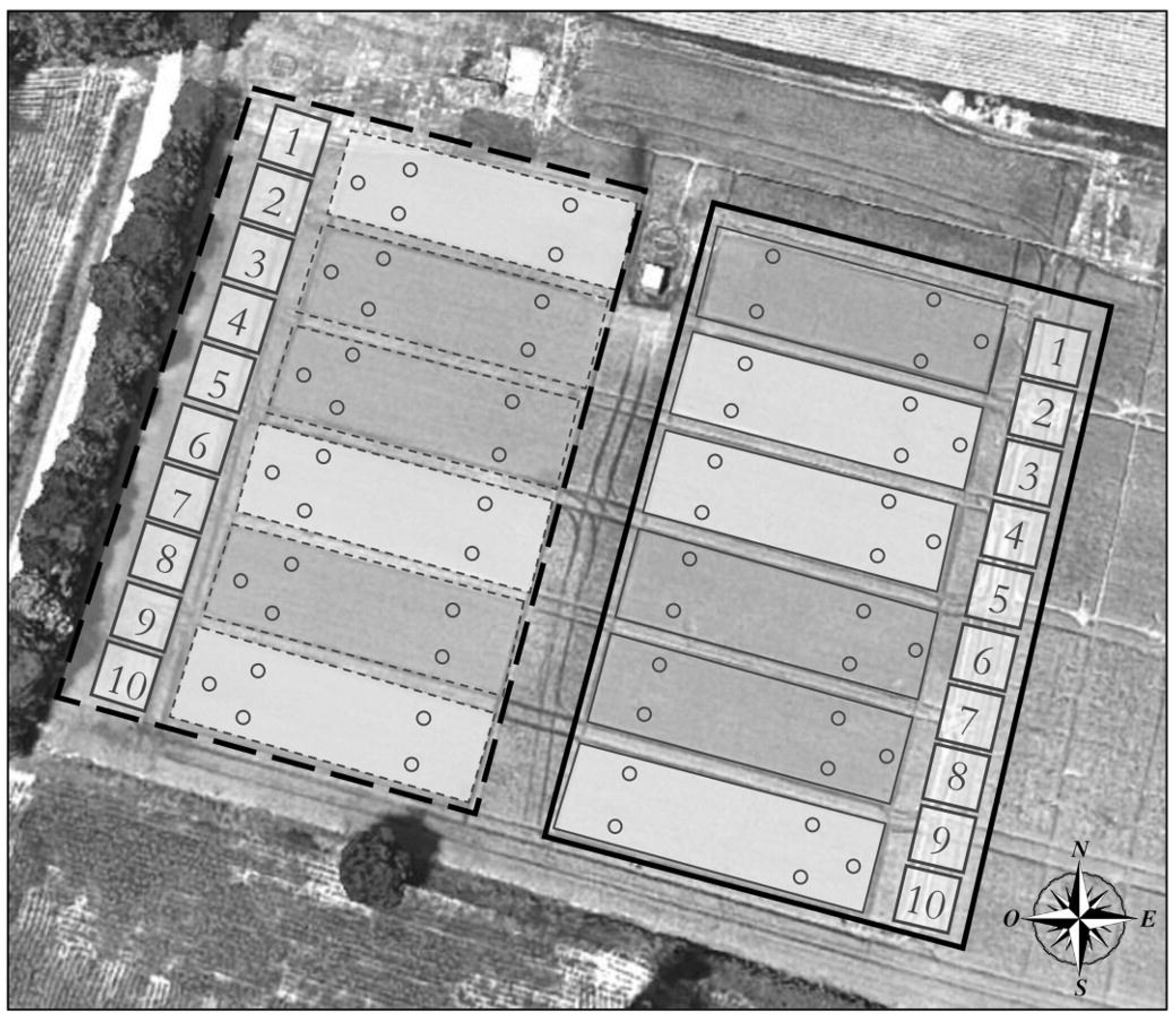
Experimental split-plot design at EFELE platform. Tillage practices determined the main plots (plowing in solid line polygons and reduced tillage in dotted line polygons) and fertilizer treatments the subplots (organic fertilization in light grey and mineral fertilization in dark grey). Circles depict the position of bait traps used for wireworm monitoring on wheat in 2017 and on maize in 2018. Numbered squares are the transect areas used to assess the effect of tillage on wireworm vertical migration on maize in 2018 and changes in its population densities between 2017 and 2020.

#### Monitoring transects to assess the effect of tillage

To allow an intensive monitoring of wireworm populations without disturbing the ground in subplots, two monitoring transects (including 10 zones spaced six meters apart) were set up at the northwest border of the reduced tillage block and at the southeast border of the conventional tillage block (Fig. 1).

### Edaphic conditions

#### Soil organic carbon content redistribution

The vertical distribution of soil organic carbon was monitored in 2012 and 2016. Soil samples were collected in the top layer (0-15 cm depth) and the bottom layer (15-25 cm depth). For each plot, soil sampling was performed in six georeferenced points and the samples were mixed to create a composite sample (3 replicates for each tillage/fertilization modality). The same sampling points were used in 2012 and 2016. Organic carbon content was determined by dry combustion using an elemental analyzer (THERMOFISHER Flash 2000 ref: EA112) according to NF ISO 10694. To assess the evolution of soil organic carbon content in the upper and bottom soil layer, the variable Delta_C was calculated as follow: Delta_C= (Soil carbon content in 2016) – (Soil carbon content in 2012). Soil bulk density and aggregate stability

#### Soil physical measurements (stability and density)

Soil physical measurements (stability and density) were carried out in May 2017, after five years of experimentation. Sampling depths were chosen to reflect the working depth of the harrow used in both tillage modalities.

For soil bulk density assessment, samples were collected at 2 depths of 0-5 cm and 10-15 cm. Five undisturbed soil samples per plot were taken (30 replicates for each tillage modality) to determine bulk density (BD) using a 15 cm manual core sampler with an internal diameter of 8 cm. The samples were weighed and dried at 105 °C for 48 h to determine gravimetric water content and BD.

For aggregate stability assessment, sampling was carried out at 2 depths, 0-10 cm and 10-20 cm. At each depth, one composite sample (made of 5 sub-samples taken with a spade) per plot was stored in a plastic box for analysis. Aggregate stability was measured on air-dried and 3-5 mm calibrated aggregates according to the standardized method ISO/FDIS 1030 (2012) derived from Le Bissonnais (1996) and Le Bissonnais and Arrouays (1997). Three disruptive tests that used different wetting conditions and energies were performed: fast wetting, slow wetting and wet stirring, which correspond respectively to a heavy storm on a dry soil, a gentle rain on a wet soil and a raindrop impact on a wet soil. The mass proportion of each fraction size of stable aggregates was calculated, and the results were expressed as a mean weight diameter index (MWD) corresponding to the sum of the mass fraction remaining on each sieve multiplied by the mean inter-sieve size. MWD was calculated for each treatment (MWD FW, MWD MB and MWD SW, respectively, for fast wetting, mechanical breakdown and slow wetting). High MWD values indicate high aggregate stability.

#### Soil moisture dynamic

Soil moisture was monitored in 2018 on maize crop between the beginning of March and the end of November in each tillage treatment, to assess how soil moisture at 40 cm depth responds to rainfall and air temperature. Soil moisture (volumetric water content, VWC) was monitored with TDR sensors of the manufacturer Trase set up once at 40 cm depth. Climatic data (daily mean air temperature and daily rainfall) were obtained from the INRAE CLIMATIK platform (https://intranet.inra.fr/climatik/, in French) managed by the AgroClim laboratory of Avignon, France.

#### Wireworm monitoring

We monitored wireworm densities using two methods: (i) soil sampling and sorting to assess changes in densities over the years, and (ii) baiting (Chabert and Blot, 1992) to assess the spatial distribution in the split-plot trial and to monitor the vertical migration of larvae in transects.

#### Temporal monitoring over the years

Every June between 2017 and 2020, two soil samples of a standard volume of eight liters (a cube of 20 x 20 x 20 cm^3^) were collected randomly in each zone of each transect described above (20 replicates for each tillage modality).

#### Temporal monitoring over seasons

Assessment of vertical migration of larvae during the cropping season was carried out according to the standardized monitoring protocol of wireworm activity in the topsoil drafted as part of the European project C-IPM ElatPro (Wechselberger et al., 2019). From March to the end of October, in 2018 when the plots were grown with maize, 19 sampling sessions were carried out for a total of 920 bait trap samples collected in the transects described above.

#### Monitoring and interpolation of the spatial distribution of wireworm densities

Spatial distribution of larvae was monitored in the subplots of the split-plot trial in 2017 on wheat and in 2018 on maize. Each year, trapping was carried out at three periods differing in terms of wireworm feeding/activity patterns. The first trapping session was carried out in spring (May-June) when larvae are supposed to be active. The second was carried out in mid-summer (late July - early August) when dry pedo-climatic conditions force wireworm to sink deep into the ground and cease all activity. The last one was carried out in autumn (October in 2017 and November in 2018), after juvenile stages had hatched and the favourable pedo-climatic conditions had returned. At each sampling period, five bait traps were regularly distributed in each plot (see Fig.1) and were set for 11 days. A total of 180 bait traps were collected in each tillage modality. Collected data were used to map the spatial distribution of wireworms in the plots with the Inverse Distance Weighting (IDW) Shepard’s interpolation (Shepard, 1968). The IDW method assigns to each unobserved point the arithmetic mean of the set observed point values weighted by the inverse of their distance from the unobserved point, raised to the power p, an unknown parameter to be estimated. Calculations were carried out using the R package gstat (R Core Team, 2024).

#### Species identification

Collected wireworms were counted and identified to species level using molecular barcoding (Folmer et al. 1994) or multiplex PCR (Staudacher et al. 2010; Mahéo et al. 2020).

#### Size distribution

Wireworms were measured using a magnifying stereomicroscope coupled with an image analysis software (Microvision Instruments, Histolab v8.1.0, Evry, France) and the proportion of young larvae in each tillage treatment was calculated. We defined as “young larvae” wireworms less than 8.5mm long (Sufyan et al., 2014).

### Statistical analysis

All statistical analyses were performed using R software (R Core Team, 2024). The effects of tillage practices and fertilization on wireworm abundances were assessed by fitting generalized linear mixed models (Bolker et al., 2009), using the ‘glmer’ function of the package ‘lme4’ (Bates et al., 2015) for count data (distribution: Poisson, link: log). Tillage and fertilization were included as fixed factors, time as a covariate and their interaction as a fixed factor. The repetition was included as a random factor. The effects of tillage, fertilization, time and their interactions were tested with a Wald test. The effect of tillage treatment on organic matter content and aggregate stability, were estimated using classical linear models using the ‘lm’ function of the package ‘stats’. Type-II analysis-of-variance tables were obtained using Anova function of the package ‘car’ (Fox and Weisberg, 2019) and TukeyHSD function of the package ‘stats’ to compute Tukey Significant Differences between treatments. The proportion of youngest larvae in tillage treatments were compared using the prop.test function (package ‘stats’).

## Results

### Edaphic conditions

#### Redistribution of soil organic carbon content between soil layers

Figure 2 shows how the organic carbon content (Delta_C) changed in the top (0-15 cm) and bottom (15-25 cm) soil layers between 2012 and 2016, according to soil tillage and fertilization type. In plowed plots, the same pattern is observed in both soil layers due to soil inversion. When combined with mineral fertilization (Fig.2A), soil organic carbon content tends to slightly decrease while it tends to slightly increase when plowing is combined with organic fertilization (Fig.2B). Conversely, when reduced tillage is applied, we observe an increase in soil organic matter content in the top soil layer (Fig.2C and 4D), particularly when reduced tillage is combined with organic fertilization, and a decrease in soil organic matter content in the bottom layer due to carbon mineralization.

**Fig. 2.**
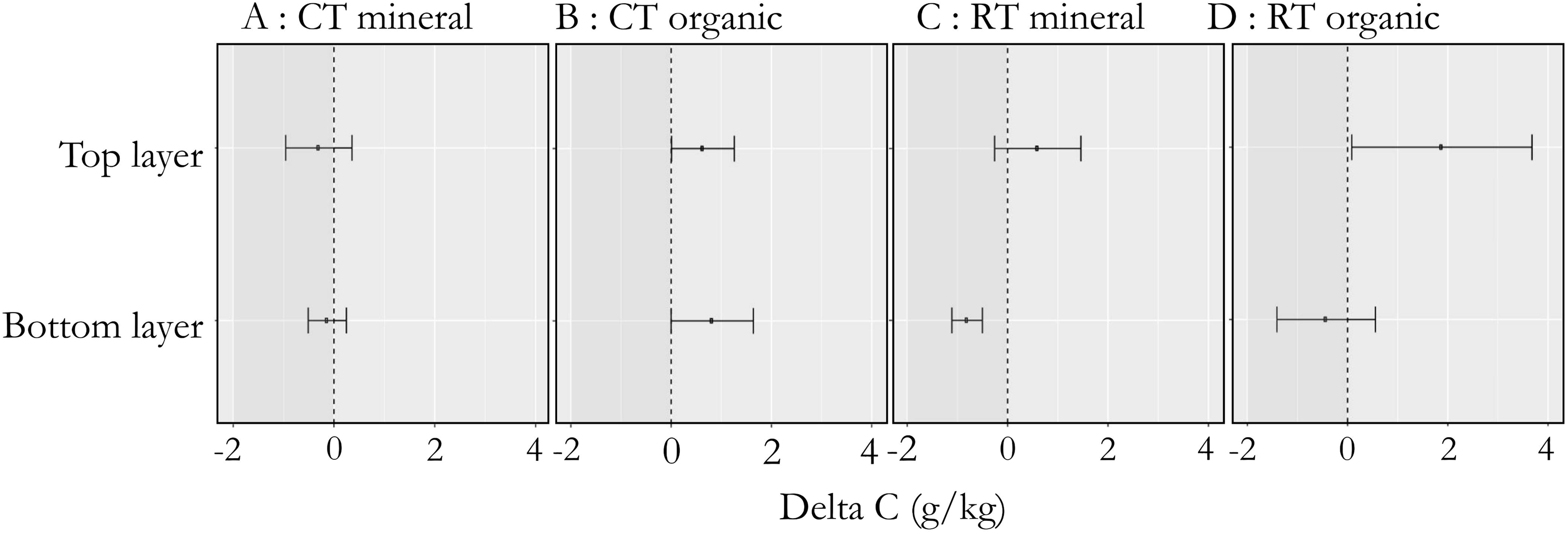
Evolution of soil organic carbon content distribution between 2012 and 2016 according to soil tillage and fertilization. CT: conventional tillage. RT: reduced tillage.

#### Soil bulk density

The vertical distribution of soil bulk density differed between tillage modalities (see Fig.3). On the top soil layer, soil bulk density was significantly lower in reduced tillage plots (ls means = 1.29) compared to plowing plots (ls means = 1.37) (χ² = 16.9, df = 1, P =0.0001). In contrast, on the bottom soil layer, soil bulk density was significantly higher in reduced tillage (ls means = 1.51) compared to plowed plots (ls means = 1.34) (χ² = 25.15, df = 1, P =0.0001). Conversely, soil bulk density was not significantly affected by fertilization type, either for the same soil layer or for the same tillage modality. Top layer soil bulk densities were very similar whether the treatment was mineral or organic in plowed plots (MF=1.36, OF=1.38), as in reduced tillage plots (MF=1.29, OF=1.28). In bottom soil layer, we observed the same similarity in soil bulk density whatever the fertilization treatment, in plowed plots (MF=1.33, OF=1.35) and in reduced tillage plots (MF=1.49, OF=1.54).

**Fig. 3.**
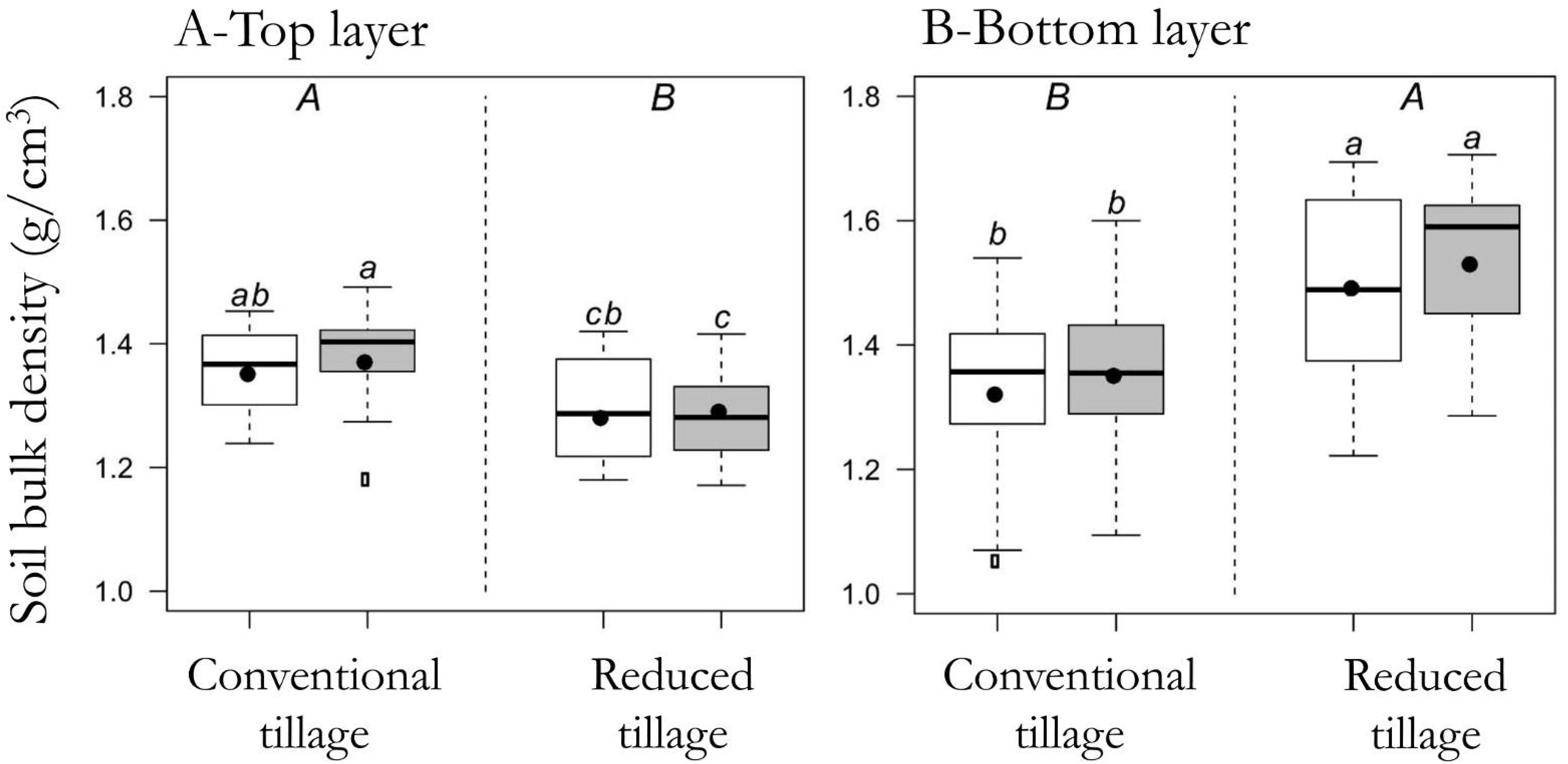
Soil bulk densities according to soil tillage and fertilization. White boxplots represent mineral fertilization (MF) and organic fertilization (OF). Uppercase letters code the significance of differences between tillage treatment and lowercase letters code the significance of differences between tillage-fertilization combinations (P-value<0.05, Ls means).

#### Aggregate stability

Figure 4 shows the mass proportion of each fraction size of stable aggregates as a mean weight diameter index (MWD). In the top soil layer, using fast wetting procedure, mean size of stable aggregates was significantly higher in reduced tillage plots (MWD=0.45 mm) than in plowed plots (MWD=0.34 mm) (χ² = 7.72, df = 1, P =0.005). Using slow wetting procedure, mean size of stable aggregates was also significantly higher in reduced tillage plots (MWD=0.72 mm) than in plowed plots (MWD=0.47 mm) (χ²= 50.6, df = 1, P =0.0001). Differences between Tillage*Fertilization treatments are also significant notably between plowing*mineral fertilization treatment (ls means = 0.43 mm) and reduced tillage*organic fertilization (ls means = 0.82 mm). Using wet stirring procedure, mean size of stable aggregates was also significantly higher in reduced tillage plots (MWD=1.51 mm) than in plowed plots (MWD=1.30 mm) (χ² = 7.72, df = 1, P =0.005) especially when combined with organic fertilization (MWD=1.81 mm). In the bottom soil layer, mean size of stable aggregates was also significantly higher in reduced tillage plots than in plowed plots for slow wetting (CT = 0.55 mm; RT = 0.64 mm) (χ² = 9.41, df = 1, P =0.002) and wet stirring (CT = 1.42 mm; RT = 1.61 mm) (χ² = 11.92, df = 1, P =0.0005).

**Fig. 4.**
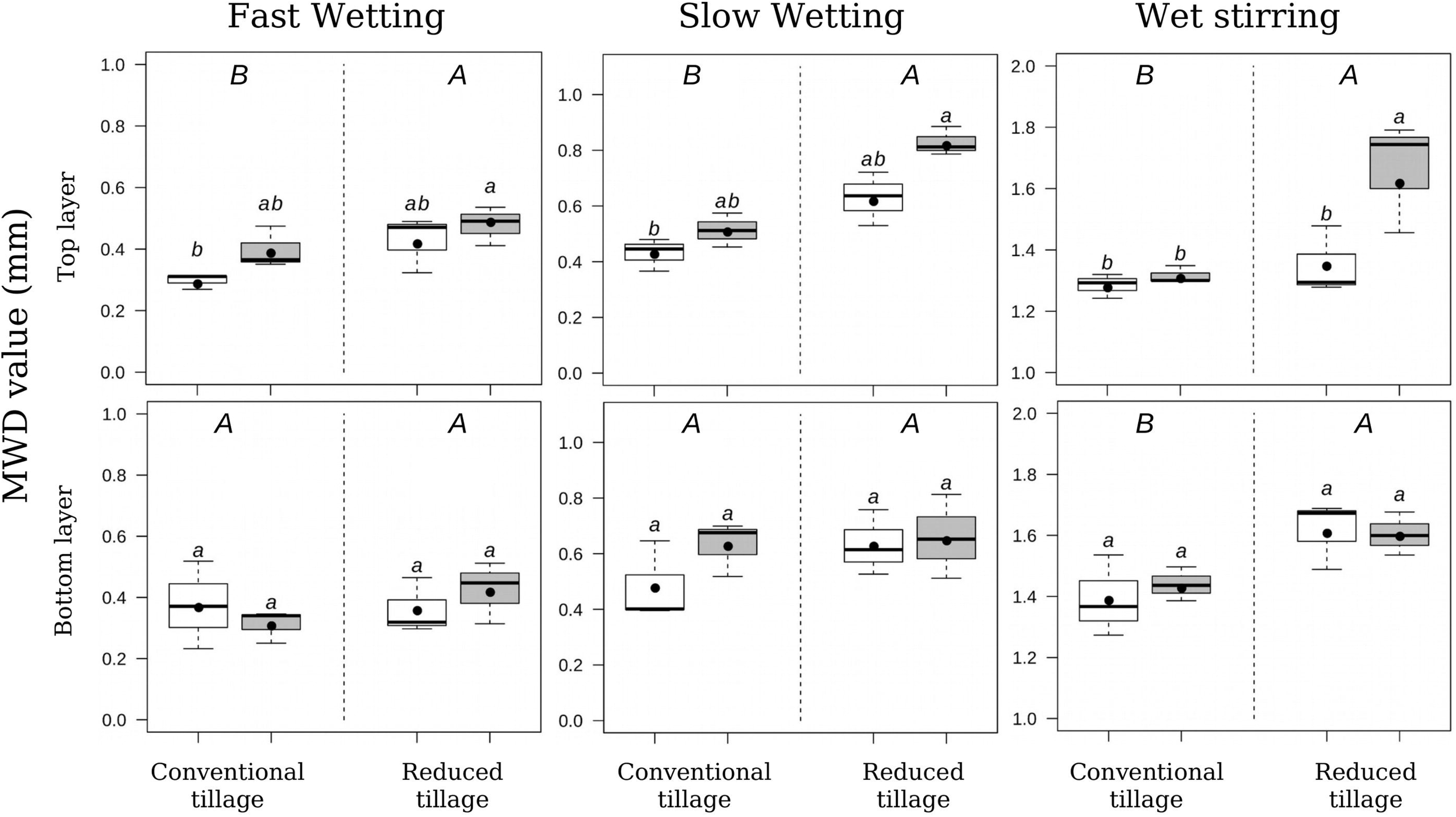
Aggregate stability according to soil tillage and fertilization. White boxplots represent ammonium nitrate fertilization and grey boxplots cattle manure. Uppercase letters code the significance of differences between tillage treatment and lowercase letters code the significance of differences between tillage-fertilization combinations (P-value<0.05, LS Means).

#### Effect of reduced tillage on soil moisture stability during maize cropping season

As shown on Figure 5, the variation of soil moisture over the maize growing season in response to changes in mean air temperature and precipitation was sharper in plowed plots (red dotted line) than in unplowed plots (green dotted line). For instance, at the beginning of June, in the absence of rainfall and with air temperature rising (Fig.4, Circle A), soil moisture abruptly decreased to less than 30% in CT while in RT it remained continuously above 30%. On the contrary, after heavy rainfalls at the beginning of autumn (Fig.4, Circle C), the soil moisture sharply increased from 15% to 25% in plowed plots while it rose only from 18% to 23% in unplowed plots. As a consequence, soil moisture variation was more buffered in the unplowed plots than in the plowed plots. Indeed, in mid-summer when temperatures were high and rainfall low (Fig.4, Circle B), the minimum soil moisture in unplowed plots was 17 % while it dropped to 13% in plowed plots.

**Fig. 5.**
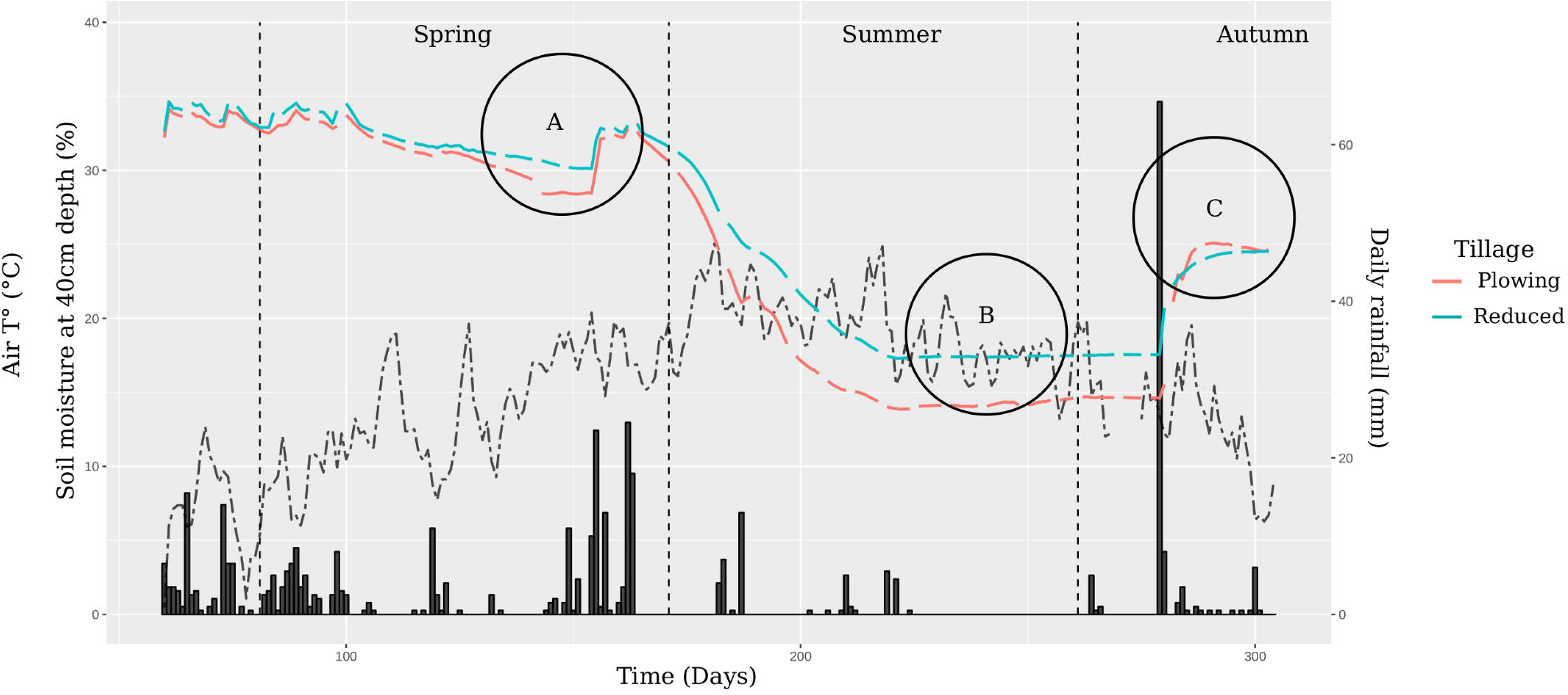
Variation of soil moisture according to weather and soil tillage. The black dotted line depicts mean daily temperature (°C), black bars depict daily rainfall (mm) and the red and blue dotted lines depict daily soil moisture (%) at 40 cm depth under plowing and reduced tillage, respectively.

### Wireworm populations

#### Species identification

In all treatments and all years, the most prevalent *Agriotes* species were, *Agriotes sputator* (67%), *Agriotes obscurus* (20 %) and *Agriotes lineatus* (13%) confirming the findings of a previous survey (Larroudé, 2015) carried outt in the northern part of France.

#### Temporal monitoring over the years in transects

Overall, we collected, between 2017 and 2020, 351 larvae in 160 soil samplings (i.e., an average of 2.19 wireworms per soil sampling). Figure 6 shows the variation in the average larval densities as a function of year and tillage treatments. The average number of wireworms was significantly higher in unplowed plots (ls means = 3.67) than in plowed plots (ls means = 0.71) (χ² = 93.5, df = 1, P =0.0001) and tended to increase in both treatments, particularly between 2019 and 2020 (from 0.70 larvae to 1.05 in plowed plots and from 3.55 to 4.45 in unplowed plots).

**Fig. 6.**
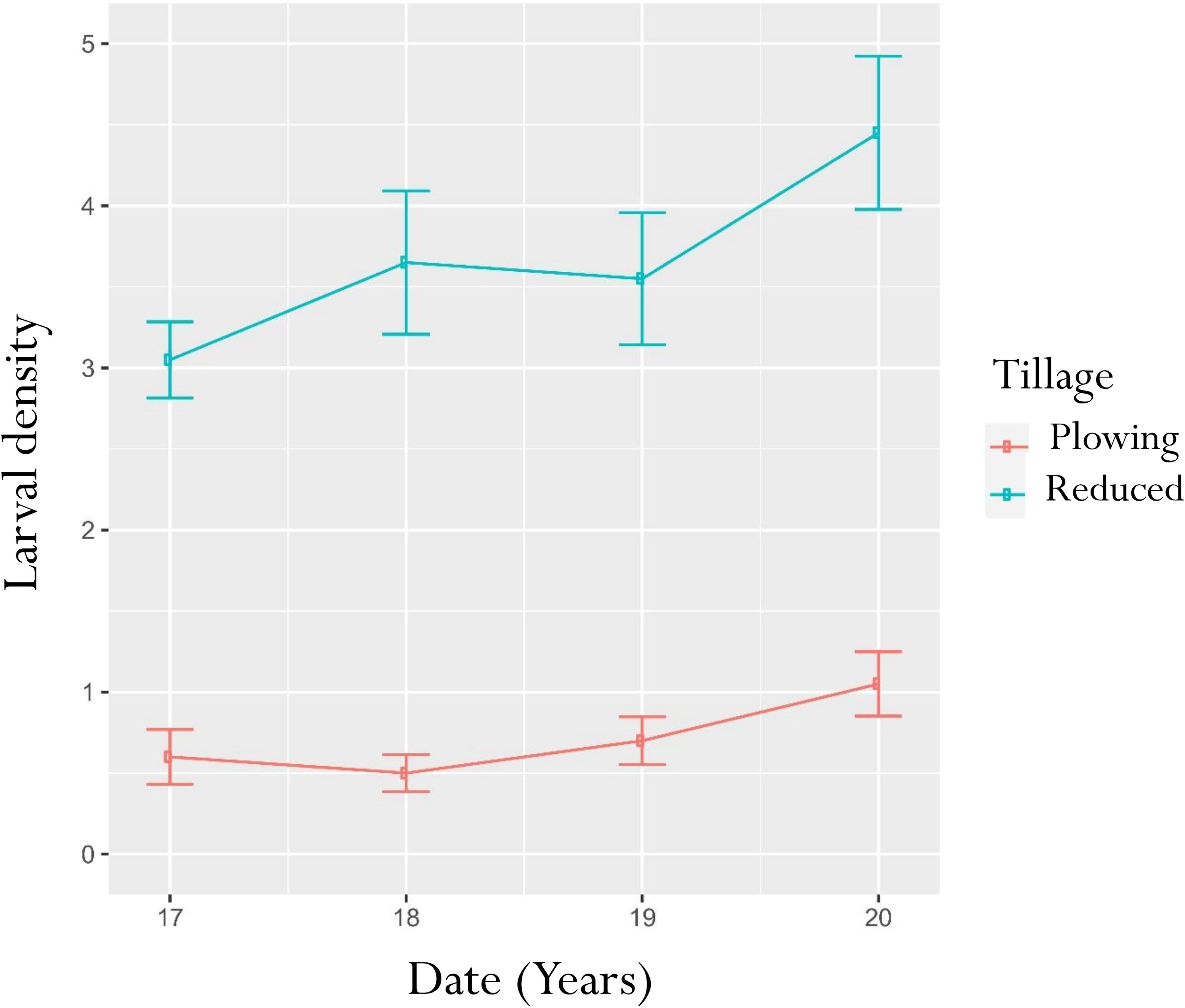
Changes in average wireworm densities in soil samplings as a function of tillage treatments between 2017 and 2020. Mean number of larvae per 20 l soil sample under plowing (red line) and reduced tillage (blue line).

#### Seasonal changes in wireworm captures

Figure 7 shows the changes in the number of wireworms captures and in soil moisture at 40 cm depth over 19 trapping periods from March to the end of October 2018. A total of 362 wireworms were trapped in 923 bait traps, for an average of 0.39 larvae per bait trap. Data confirmed higher abundance in the unplowed zone, where we caught 292 larvae for an average of 0.63 larvae per bait trap whereas we caught only 70 larvae for an average of 0.15 larvae per bait trap in the plowed zone. Pairwise comparisons showed significant differences in the average number of captures between treatments at almost all sampling dates. Data did not indicate a clear relation between the number of captures and the soil moisture but the highest infestation level was observed under the driest conditions.

**Fig. 7.**
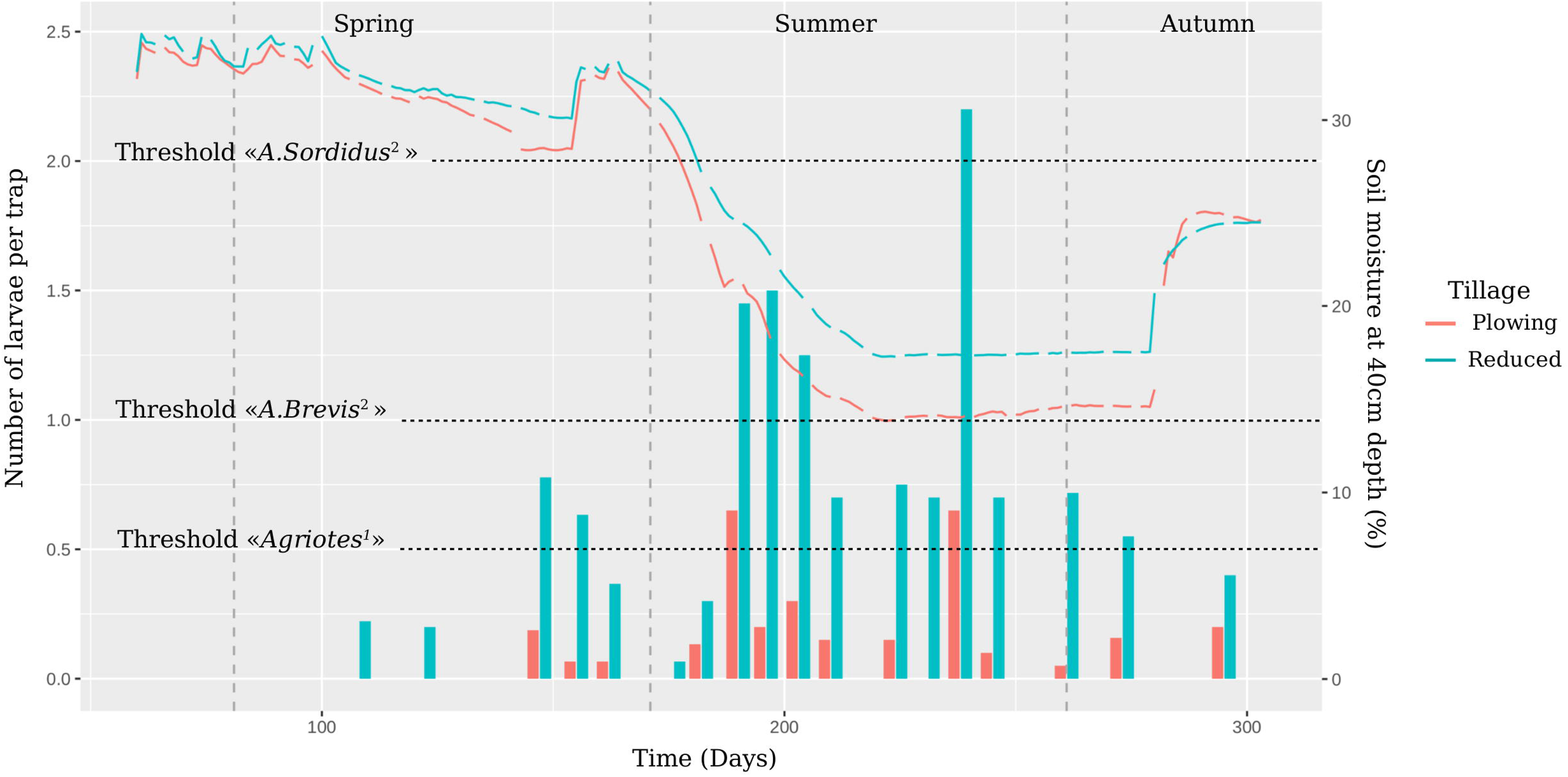
Seasonal dynamics of trapped wireworm abundance. Bars represent the mean number of wireworms captured per trap at each sampling date (between 16 and 40 replicates per date) and colored dotted lines represent daily soil moisture (%) at 40cm depth as a function of tillage treatment. Black dotted lines represent IPM thresholds proposed by (1) Chabert at al.1993 and (2) Furlan, 2014.

#### Spatial distribution of wireworm captures in subplots

A total of 137 wireworms were trapped in the 360 bait traps, 61 in 2017 on winter wheat and 76 in 2018 on maize. Figure 8 shows the inferred spatial distribution of wireworm infestation levels in plots and highlights the aggregated distribution of wireworms and an intensification of infested foci between 2017 and 2018.

**Fig. 8.**
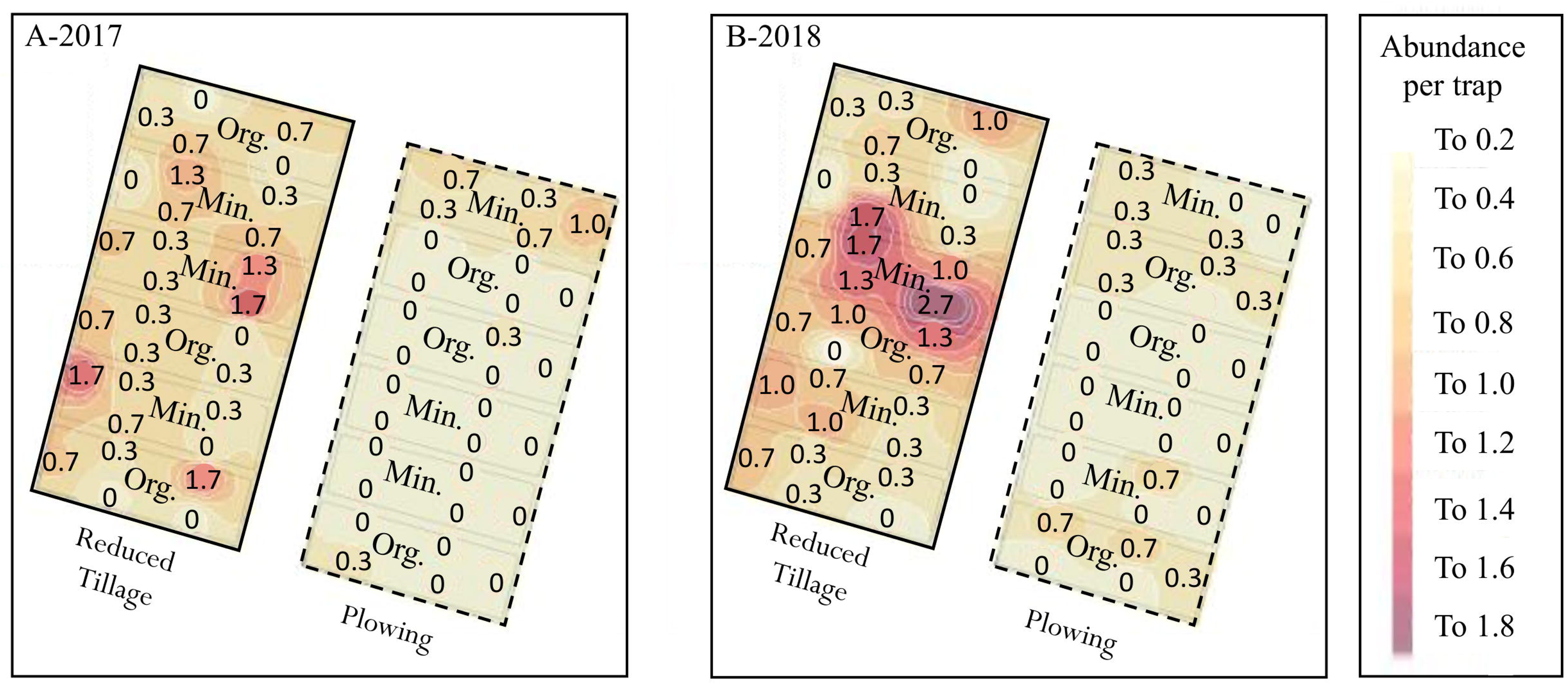
Spatial distribution and infestation levels of wireworms using Inverse Distance Weighting Interpolation. Numbers represent the abundance of wireworms at each trap location averaged over the three sampling dates (three replicates).

#### Wireworm infestation levels in subplots according to soil tillage and fertilization

Figure 9 shows the average number of wireworms captured per bait trap as a function of soil treatment (tillage and fertilization). As with transect sampling, wireworm infestation levels were significantly higher in unplowed plots than in plowed plots. In 2017, only 10 wireworms were trapped in plowed plots (lsmean = 0.09 larvae per trap) compared to 50 in unplowed plots (lsmean = 0.53 larvae per trap) (χ² = 18.3, df = 1, P =0.0001). In 2018, 14 were trapped in plowed plots (lsmean = 0.15 larvae per trap) whereas 62 larvae were trapped in plowed plots (lsmean = 0.67 larvae per trap) (χ² = 21.1, df = 1, P =0.0001). In contrast, the average number of wireworms collected in subplots did not seem influenced by fertilization. Although wireworm infestation levels in 2017 were significantly higher in plots with mineral fertilization (lsmean = 0.36 larvae per trap) compared to plots with organic fertilization (lsmean = 0.14 larvae per trap) (χ²= 5.04, df = 1, P =0.02) whatever the tillage treatment, the interaction between tillage and fertilization was not significant (χ² = 1.47, df = 1, P =0.22), in 2018, fertilization treatments did not differ anymore.

**Fig. 9.**
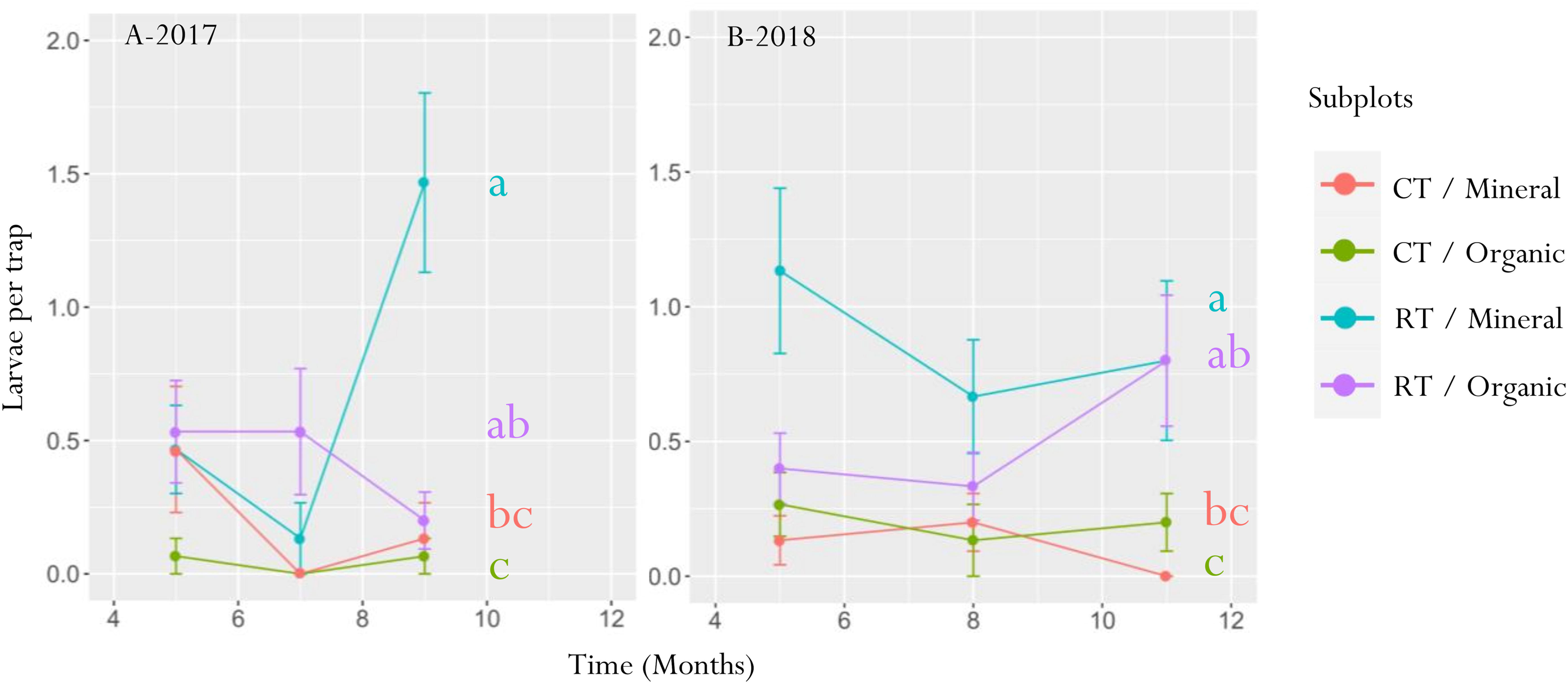
Wireworm infestation levels in subplots according to soil tillage and fertilization.

#### Size distribution in bait traps

Figure 10 shows the size distribution of the *Agriotes* larvae in relation to the tillage treatment collected in soil samples (Fig.10.A) or captured in bait traps (Fig. 10B). A higher proportion of young larvae (less than 8.5 mm long) were caught in plowed plots than in unplowed plots, whatever the method used. Regarding larvae collected in soil samples during the four years of monitoring, only 9% were considered as young in the reduced tillage zone compared to 35% in the plowed zone. The X² value (18.21 with 1 degree of freedom) from the prop.test function (package Stats) gives a significative p-value (p=0.001). Regarding larvae caught using bait traps in transects and in the split-plot trial, 21% were considered as young in plowing compared to 46% in reduced tillage. (χ² = 13.32, df = 1, P =0.001).

**Fig. 10.**
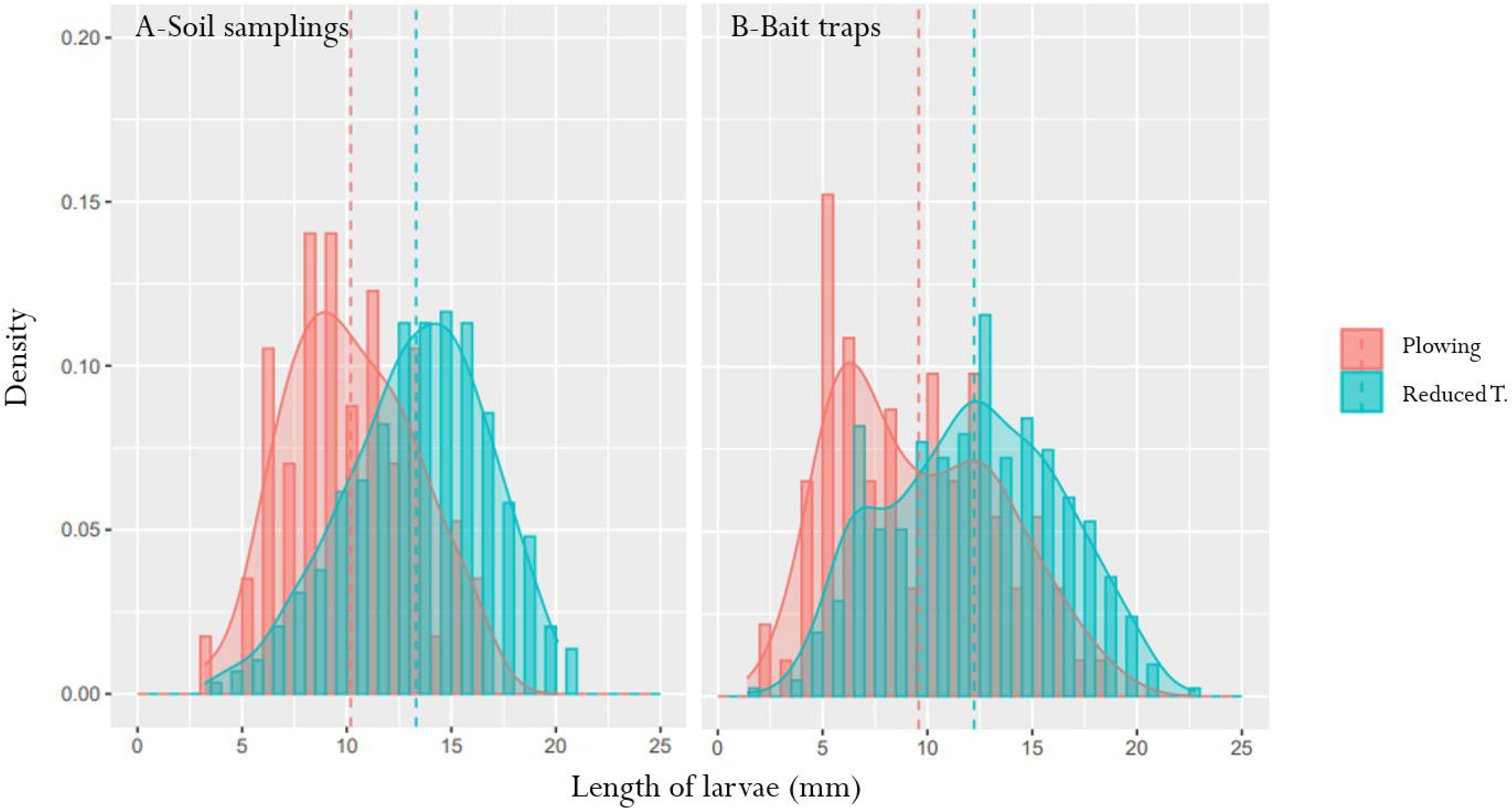
Distribution of *Agriotes* larval sizes in bait traps depending on the tillage treatment.

## Discussion

In this study, we compared a range of abiotic parameters (vertical distribution of soil organic carbon, soil bulk density, aggregate stability, soil moisture variability) and biotic features (wireworm abundance and spatial distribution) after five years of reduced tillage or plowing, the minimum duration required for system stabilization (Bai et al., 2018). With the exception of soil bulk density, long term reduced tillage tended to improve soil physical properties but in the same time to increase infestation level by wireworms.

### Soil organic matter distribution

Franzluebbers (2002) argued that a clear stratification of soil organic matter content with depth indicates soil quality and a well-functioning of soil ecosystem, because surface organic matter is essential for controlling erosion, facilitating water infiltration, and conserving nutrients. In our study, we observed an increase in soil stratification in terms of organic matter content, particularly when reduced tillage was combined with organic fertilization. This was due to the accumulation of organic matter in the upper soil layer and its subsequent decrease through carbon mineralization in the lower layer. Under conventional tillage, the organic matter content was not vertically stratified due to soil inversion.

### Soil bulk density

Soil bulk density is a primary indicator of soil compaction and structural health. For example, Bruand et al., (1996) demonstrated the importance of SBD for water holding which can quadruple as SBD decreases from 1,8 g/cm^3^ to 1,2 g/cm^3^. On the contrary, high SBD values indicate a poor soil structure and a low macroporosity that can negatively affect plant root growth but also increase the volume of medium pores (Rasmussen, 1999). In our study, one of the most noticeable effects of reduced tillage on edaphic conditions is an increase in soil bulk density in the bottom soil layer. As previously showed by (López-Fando and Pardo, 2011; Rasmussen, 1999), in plowed plots (see figure 2), SBD values in the top and the bottom soil layers were similar, which may essentially stem from homogenization induced by mouldboard plowing. In reduced tillage plots, the rotary harrow induced low SBD values on the surface while soil was much more compact at a depth below 10 cm under the tilled soil layer. This compaction may be due to the massing of the silty soil, which had not been tilled up for five years, and to the passing of the manure spreader. Our results are in line with Pöhlitz et al., (2024), who found that cultivator tillage (RT) led to a significant increase in soil bulk density, reaching approximately 1.50 g cm□³ in the second year, compared to the ∼1.30 g cm□³ under conventional plough tillage (CT).

### Aggregate stability

Finally, in line with the results reported by (Bottinelli et al., 2017), aggregate stability in the top soil layer where the organic carbon is concentrated was improved under reduced tillage, especially when combined with organic fertilization. Aggregate stability is associated to several soil properties including erodibility, carbon storage or root penetration (Bottinelli et al., 2017) and contributes to the maintenance of soil porosity as well as a good water infiltration. Thus, our results outline the beneficial effects of reduced tillage on erosion reduction (Palm et al., 2014).

### Soil moisture stability

It is generally accepted that reduced tillage improves the efficiency of rainwater use by augmenting soil water storage (Benites and Castellanos-Navarrete, 2003; Findlater, 2013; Kassam et al., 2014). The observed dynamics of soil water content at 40 cm depth illustrates the buffering of soil moisture in reduced tillage plots compared to plowed plots. As previously showed (Belmekki et al., 2014), soil moisture responded in our study more sharply to temperature and rainfall changes in plowed plots than in reduced tillage plots. For example, in early June, rising temperatures and absence of rainfall caused a rapid drop in soil moisture below 30% in plowed plots, while it constantly stayed above 30% in reduced tillage plots. After heavy autumn rain, soil moisture increased more sharply in plowed plots (from 15% to 25%) than in unplowed plots (from 18% to 23%). Overall, soil moisture in reduced tillage showed fewer extreme fluctuations.

### Wireworm populations

Wireworm populations could be directly affected by plowing. Our study confirmed the findings of previous studies (Le Cointe et al., 2023; Salt and Hollick, 1949; Seal et al., 1992; Wechselberger et al., 2019), which showed that plowing was effective in limiting wireworm populations. Indeed, the number of wireworms collected in transect samples in June was consistently divided by about five in plowed plots compared to reduced tillage plots over the four years of monitoring. The diachronic study carried out in 2018 in transects using bait traps confirmed a strong reduction of wireworm abundance in the plowed zone throughout the growing season compared to the unplowed zone. Until now, the impact of tillage reduction was supposed to result from the destruction of eggs and young larvae (Fox, 1961; Furlan, 2005; Ritter and Richter, 2013). It has been demonstrated that the decrease in the number of wireworms due to repeated soil disturbance after grassland cultivation leads to a significant decline in the number of young larvae. In 1949, Salt and Hollick conducted a five-year experiment which revealed that the decline in wireworms was accompanied by a significant change in the distribution of larval sizes, as evidenced by a reduction in the number of young larvae. It is also currently acknowledged that the lack of soil cover, especially in springtime, might limit oviposition (Gough and Evans, 1942). However, the higher proportion of young instars in plowed plots than in unplowed plots suggests that the absence of soil cover did not prevent oviposition in our study. Therefore, we assume that the infestation reduction we observed in plowed plots was due more to direct plow injury or increased vulnerability to predation or desiccation than to the prevention of oviposition.

### Soil moisture

Soil moisture is known to influence wireworm population dynamics. Lefko et al., (1998), emphasize the significant role of soil moisture in wireworm survival and spatial distribution. They suggest that soil moisture could indicate areas where wireworms are more prevalent and guide scouting efforts within a field. In laboratory experiments, (Campbell, 1937) demonstrated that larvae die if they remain in dry soil for too long and stop their activity if they have to stay in water saturated soil. In consequence of a drying out soil, wireworms can also migrate in the bottom soil layer to reach humidity (Furlan, 1998; Jung et al., 2014). Soil moisture can also affect the feeding behaviour of wireworms. (Samoylova and Tiunov, 2017) demonstrated that *Agriotes obscurus* larvae in the Russian steppe switch from a herbivorous regime in winter, when the steppe is wet, to a carnivorous regime in summer, when the steppe is dry. It is also commonly established that wireworms have two seasonal peaks of abundance in the upper soil layer, the first from March to May and the second from September to October (Furlan, 1998; Gratwick M, 1989) which are thought to depend on soil moisture and temperature (Jung et al., 2014; Lafrance, 1968). In short, it is said that wireworms seek shelter in the lower soil layer during the dry summer months. However, the seasonal monitoring performed in this study, which aimed to describe this vertical migration of larvae, shows the opposite to be true. The first seasonal population peaks were observed when soil moisture was in depletion and the highest infestation level was observed under the driest conditions. These results could be explained by wireworms behavior be driven by water and not only by food.

### Organic matter

Finally, we also highlight the redistribution of organic matter in the upper soil layer that could be profitable to wireworms population (Furlan et al., 2016; Kozina et al., 2015; Poggi et al., 2018). Several studies have examined the relationship between soil organic matter content and wireworm populations. Salt and Hollick (1946) reported a link between the distribution of wireworms and the level of organic matter in the soil. This was based on the theory that wireworms feed on organic matter. But Traugott et al. (2008) showed, using isotope analysis, that wireworms consume only a negligible amount of organic matter. Still, the content of organic matter in the soil may influence its suitability for wireworms by affecting soil structure. This can hinder or facilitate the movement of wireworms and the detection of food via volatile compounds released in the rhizosphere (Gfeller et al., 2013). Several studies have examined the relationship between the content of soil organic matter and wireworm damage. Findings suggest that soil organic matter higher levels are linked to greater wireworm damage. For example, Poggi et al. (2018) found that soil with organic matter content above 5.5% should be considered as risky for maize seedlings.

### Wireworms ecology in conservation agriculture systems

This study shows that, while improved physical properties are indeed observed, this comes at the cost of an increased wireworm population. Therefore, a compromise must be found between the different ecosystem services and disservices related to soil. Stopping plowing is usually the first step towards conservation agriculture (CA) which combines three principles (1) limitation of soil disturbance, (2) its permanent cover and (3) crop diversification. Despite the slightly provocative nature of this article’s title, we would like to conclude by reiterating that, since it is now recognized that stopping plowing improves soil health and increases biodiversity, including earthworms, it should not be surprising that it also increases the number of organisms considered to be pests, such as wireworms. However, this should not be considered a threat in terms of crop damage. Indeed, despite potentially high infestation levels in CA systems, economic damage to maize crop is rarely observed (Furlan et al., 2021). If stopping plowing is usually the first step toward conservation agriculture, it is an incomplete implementation of this. Soil permanent cover should considered as utmost importance as its allow to deal with wireworms without damage by supplying them fresh organic matter on which they can feed (Le Cointe et al., 2023; Sonnemann et al., 2012). Finally, in our study, the experimental crop rotation was quite simplist, consisting in a two-year succession of maize and winter wheat. And crop diversification can affect directly wireworm populations by breaking the continuity of resources over time and space, or indirectly, by favouring natural enemies.

## Conclusion

In our study, we investigated whether stopping plowing would have a synergistic effect on improving edaphic conditions and reducing pest pressure. Our results show that, while improved physical properties are indeed observed, this comes at the cost of an increased wireworm population. Our study highlights the importance of adopting a holistic approach when designing sustainable cropping systems, as well as considering the services and dis-services they provide.

## Acknowledgements

This study was carried out as a part of the project STARTAUP “Design of alternative strategies for controlling wireworm damage in maize crops” with financial support from The Foundation for Research on Biodiversity (FRB, project neonic-33) as part of the call on “Sustainable crop protection without neonicotinoids: improving the emergent and opening innovative perspectives”. The EFELE field experiment forms part of the SOERE-PRO (network of long-term experiments dedicated to the study of impacts of organic waste product recycling) certified by ALLENVI (Alliance Nationale de Recherche pour l’Environnement) and integrated as a service of the ‘‘Investissement d’Avenir’’ infrastructure AnaEE-France, overseen by the French National Research Agency (ANR-11-INBS-0001). The authors are grateful to Philippe Le Roy and Florian Gaillard for their contribution to field experiments. RLC and SP thank partners of the ElatPro project (ERA-NET C-IPM, 2016-2019, grant no.:618110) for meaningful discussions.

## Notes

### Competing Interest Statement

The authors have declared no competing interest.

